# CollapsedChrom: resolving the assembly of collapsed chromosomal segments in polyploid genomes of the model grass genus *Brachypodium*

**DOI:** 10.64898/2026.03.07.710290

**Authors:** Wenjie Mu, Jianquan Liu, Pilar Catalán

## Abstract

Polyploidization plays a fundamental role in plant evolution and crop domestication. However, due to the high similarity of genomic sequences between some homologous or homeologous chromosomes, the assembly of some polyploid genomes is extremely difficult, frequently resulting in erroneous assemblies, such as sequence chimeras and sequence collapse. The genus *Brachypodium* is an important model system for the study of polyploidy in grasses. However, high-quality reference genomes are still lacking for its complex polyploid perennial species. In this study, we developed a bioinformatic pipeline for the accurate assembly of high-quality reference genomes at the chromosomal level for two representative perennial *Brachypodium* species with conflicting collapsed segments, the allotetraploid *B. phoenicoides* (2n = 4x = 28) and the autohexaploid *B. boissieri* (2n = 6x = 48). We developed an innovative methodology (CollapsedChrom) that uses depth-of-read profiling and relies on prior karyotypic information to systematically detect and rescue collapsed regions. This depth-sensitive curation strategy successfully recovered 328.9 Mb and 195.8 Mb of previously collapsed sequences in the genomes of *B. phoenicoides* and *B. boissieri*, respectively. Comprehensive quality assessments demonstrated the high quality of our final assemblies. Our chromosomal-level assemblies fully capture the genomic architectures of these species. These robust genomic resources overcome long-standing challenges in polyploid assembly and provide an essential foundation for future research on the evolutionary dynamics, subgenomic interactions, and functional biology of complex polyploid plant genomes.

## Introduction

Polyploidization has occurred repeatedly across the tree of life and is particularly prevalent in plants^1^. Many polyploid species are important crops, such as wheat, oat, strawberry, peanut, sugarcane, and potato^1,2^. Furthermore, even if their diploid progenitors remain undiscovered or have become extinct, the polyploid wild relatives of these crucial species could still harbor large amounts of genetic information that was lost during crop domestication due to domestication bottlenecks^3–5^. This genetic reservoir could facilitate further crop improvement^6^. However, polyploidy extends beyond cultivated species and their wild relatives, encompassing approximately 50% (30% to >70%) of species diversity in terrestrial plants^7^. Polyploidy enhances plant adaptation and resilience by reconfiguring metabolic pathways and optimizing survival in challenging environments^8^, supporting the success and recurrent formation of plant polyploids throughout evolutionary time^9^.

The grass genus *Brachypodium* includes species selected as model systems for functional genomics of grasses and monocots^10,11^, and for analyzing the recurrent origins of allopolyploidy^12,13^ and the evolutionary switches from perenniality to annuality^14^. This genus exhibits exceptionally rich karyotypic diversity^15,16^, with chromosomal rearrangements resulting in various chromosome basic numbers (x = 10, 9, 8, 5), ploidy levels ranging from diploid to hexaploid, and several descending dysploidy trends, providing an ideal natural system for studying karyotype evolution, alternative polyploidization scenarios, and their evolutionary consequences. Among them, the annual allotetraploid *B. hybridum* and its two diploid progenitor species, *B. distachyon* and *B. stacei*, have been well studied^12,13,17^.

However, in the recently diverged core perennial clade, recurrent chromosomal rearrangements and multiple polyploidization events have driven an extensive radiation of species and cytotypes^16^. Among the major subgenomes (A2, E1, E2, and G) derived within this clade, with the exception of subgenome G, which contains extant diploid species with a similar karyotype, diploid species with karyotypes similar to subgenomes A2, E1 and E2, remain undiscovered or extinct^16^. Representative species include the tetraploid *Brachypodium phoenicoides* (2n = 4x = 28), thought to comprise the G and E2 ghost subgenomes, and the high-ploidy hexaploid *B. boissieri* (2n = 6x = 48), which contains three ghost A2 subgenomes with identical karyotype^16^. While these species are very crucial for decoding the mechanisms, consequences, and evolutionary adaptation of polyploidization and descendant dysploidy trends, high-quality reference genomes for them are still lacking due to their highly complex ploidy features.

The assembly of polyploid genomes is inherently challenging; their complex genomic architectures typically contain highly similar or even identical homologous sequences^18^. This similarity originates not only from homologous sequences between the two haplomes inherited from the parents but also potentially from homeologous chromosomes across subgenomes (allopolyploids) and/or haplotypes (autopolyploids). Especially when there are multiple chromosomes from closely related karyotypes, widespread homeologous exchanges^19^ are more likely to lead to frequent fragmentation and misassembly, including sequence chimerism and sequence collapse^18^. Sequence chimerism occurs when fragments from different genomic regions (such as distinct subgenomes or physically unlinked regions) are incorrectly joined during assembly to form an artificial mixed sequence^20^. Sequence collapse occurs when multiple homologous or homeologous regions are incorrectly merged into a single sequence^21^. Although current conventional assembly strategies (such as PacBio HiFi-based assembly) handle sequence chimerism through Hi-C contact information^22,23^ and manual curation, and generate highly contiguous primary assemblies (haplome) by purging haplotig duplications^24,25^ to ignore collapsed regions^26^, this practice often results in the loss of intragenomic variation. It can particularly affect genomes with high heterozygosity, which could directly interfere with subsequent variant discovery, allele-specific differential expression analysis, subgenome identification, and accurate evolutionary history reconstruction^18,27^.

To address this challenge, the present study employed PacBio HiFi sequencing, Hi-C assisted scaffolding, and the newly released C-Phasing software^23^ to perform a comprehensive genomic analysis of the genomes of *Brachypodium phoenicoides* Bpho6 and *Brachypodium boissieri* Bbois3 polyploids. We designed and implemented an innovative CollapsedChrom pipeline, based on prior knowledge of the karyotypes of these accessions ^16^, with in-depth read alignment information to accurately detect and rescue collapsed regions during the assembly process. Using this strategy, we successfully recovered 328.9 Mb and 195.8 Mb of collapsed sequences from the initial assembly of the Bpho6 and Bbois3 genomes, respectively, and constructed chromosome-level genomes that meet reference criteria. These comprehensive, high-quality reference lines not only correct structural defects in initial assemblies, but also provide powerful tools for exploring the evolutionary dynamics and functional biology of the genus *Brachypodium* and other complex polyploid plants.

## Results and Discussion

### Genome survey

Previous cytogenetic knowledge of the genomic architecture of *Brachypodium phoenicoides* Bpho6 and *Brachypodium boissieri* Bbois3 was based on the results of genome size and chromosome count analyses, which indicated a DNA content of 1,443 ± 0.019 pg/2C and 2n=28 chromosomes for the tetraploid accession Bpho6, and 3,236 ± 0.072 pg/2C and 2n=48 for the hexaploid Bbois3^16,28^. These data provide reference values for the total size of each genome. Furthermore, karyotypic analyses using comparative chromosome barcoding indicated that Bpho6 is an allotetraploid with two subgenomes having different basic chromosome numbers, subgenome G with x=9 chromosomes and subgenome E2 with x=5 chromosomes. Conversely, Bbois3 is an autohexaploid with three copies of the same karyotype A2 with x=8 chromosomes^16^. These data were extremely useful for establishing the chromosome assembly of the subgenomes in the samples under study.

Subsequently, we performed a k-mer-based genomic study using sequencing data, combining GenomeScope2 and Smudgeplot analyses^29^. GenomeScope2 modeling yielded well-fitting k-mer spectra for both species. Bpho6 exhibited four distinct coverage peaks, while Bbois3 showed six peaks, supporting tetraploid (4x) (Fig. 1A) and hexaploid (6x) (Fig. 1B) genomic architectures, respectively. Furthermore, GenomeScope2 estimated heterozygosity levels of 2.4% for Bpho6 (Fig. 1A) and 4.38% for Bbois3 (Fig. 1B). Notably, in polyploid genomes these values likely capture not only allelic divergence between two haplotypes, but also sequence divergence between homologous copies of different subgenomes (in the case of autopolyploids, between multiple homologous copies), and should therefore be interpreted as composite measures of intragenome homologous sequence divergence rather than diploid-type heterozygosity. Smudgeplot provided independent support for the inferred polyploid architectures. In Bpho6, the dominant smudge classes were AABB (0.34) and AAAB (0.21), consistent with a tetraploid genomic composition (Fig. 1C); in Bbois3, the most abundant classes were 5A1B (0.16), AB (0.12), and AAB (0.12), supporting a hexaploid-type k-mer pairing structure (Fig. 1D). Taken together, these experimental (chromosome counting, genome size, chromosome barcoding) and in silico (GenomeScope2, Smudgeplot) results provide critical background knowledge on genome size, ploidy, karyotype, and within-genome divergence, thus guiding the subsequent assessment of genome assembly and interpretation of genome structure in these two polyploid species.

**Figure 1.**
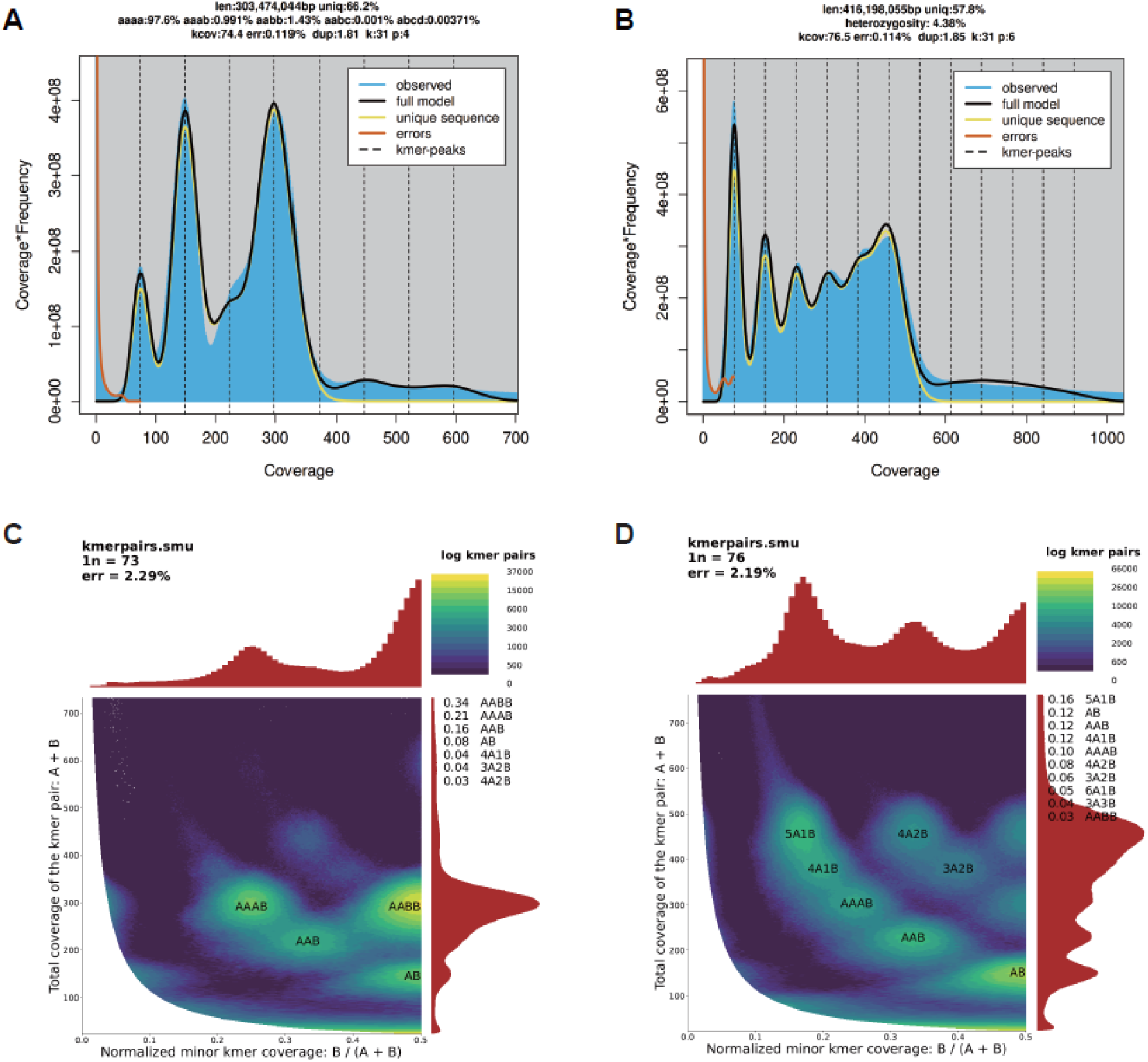
k-mer based genome survey and inference of polyploid architecture in *Brachypodium phoenicoides* Bpho6 and *B. boissieri* Bbois3. GenomeScope2 modeling of Illumina k-mer spectra for *B. phoenicoides* Bpho6 (A) and *B. boissieri* Bbois3 (B). Smudgeplot k-mer–pair analyses for Bpho6 (C) and Bbois3 (D), showing the distribution of normalized minor k-mer coverage and total k-mer–pair coverage.

### Initial assembly of the primary genome

We generated initial de novo assemblies for Bbois3 and Bpho6 using hifiasm^30^, widely designed for PacBio HiFi-based assembly. Contigs smaller than 50 kb were removed to reduce fragmentation and potential assembly artifacts. The resulting draft assembly of Bpho6 had a total length of 625,824,682 bp with an N50 contig of 26,277,391 bp, while the Bbois3 assembly totaled 1,389,706,146 bp with an N50 contig of 23,752,561 bp. According to previous karyotype studies, Bpho6 is an allotetraploid comprising the G (x=9) and E2 (x=5) subgenomes of *Brachypodium*, while Bbois3 is derived from the triplicated A2 (x=8)^16^ genome of *Brachypodium*. To assess sequence completeness and overall integrity of the primary assemblies, we performed whole-genome alignments by mapping Bpho6 to the G reference genome of *B. sylvaticum* (Bsyl-Ain1)^14^ and mapping Bbois3 to the A2 reference subgenome of *Brachypodium retusum* Bret403 (unpublished data). Although N50 contig values indicated high assembly contiguity, we unexpectedly detected apparent copy number deficits in the primary assemblies when compared to their corresponding references.

Specifically, in Bpho6, whole-genome alignment revealed a missing copy of a genomic segment homologous to *B. sylvaticum* Chr1 (two haploid copies expected in the tetraploid genome, but only one was recovered). Similarly, in Bbois3, we identified a missing copy of a segment homologous to Bret403 A2_07 (three haploid copies expected in the hexaploid genome, but only two were present) (Fig. S1). Furthermore, we used Merqury 30 to estimate the consensus quality (QV) scores of the assembly and k-mer-based completeness. In the initial primary assemblies, Bpho6 showed a low QV score of 17.10 and a completeness of 54.10%, while Bbois3 also showed a QV score of 17.19 and a completeness of 51.38%. These results indicate that, while purging haplotype duplications to generate a consensus primary assembly may improve contiguity, it could also reduce the nucleotide diversity retained in these two polyploid genomes.

### Chromosome-level genome assembly

To retain as much nucleotide diversity as possible in these Bpho6 and Bbois3 polyploid genomes, we aimed to capture all haplotype- and subgenome-specific sequences using the hifiasm unitig assembly (unitig: chromosome-level processed unitig graph without small bubbles) and performed a Hi-C guided chromosome-scale assembly with C-Phasing. After extensive manual curation of the initial scaffolding results, the chromosome-level assemblies showed a marked improvement in consensus quality and k-mer–based completeness.

Specifically, Bpho6 reached a QV of 54.71 with 98.24% completeness, whereas Bbois3 reached a QV of 46.48 with 97.63% completeness. However, despite the fact that C-Phasing combined with manual correction effectively removed chimeric assembly errors, we still detected abundant collapsed assembly errors in both the Bpho6 and Bbois3 assemblies (Figs. 2A, 2B). These collapsed regions were independently supported by Hi-C interaction heatmaps and read-alignment depth profiles. Specifically, the genome-wide distribution of read-mapping depth showed a double peak pattern in both Bpho6 and Bbois3 (Fig. S2), and chromosome-wide analyses with 100-kb windows identified regions whose median mapping depth was approximately twice that of the other regions, consistent with the collapse pattern (Figs. 2C, 2D). Notably, in Bbois3, an extreme case was observed in which an entire chromosome appeared to have collapsed, such that the expected 48 chromosomes were represented as only 47 chromosome-scale scaffolds in the initial chromosome-level assembly.

**Figure 2.**
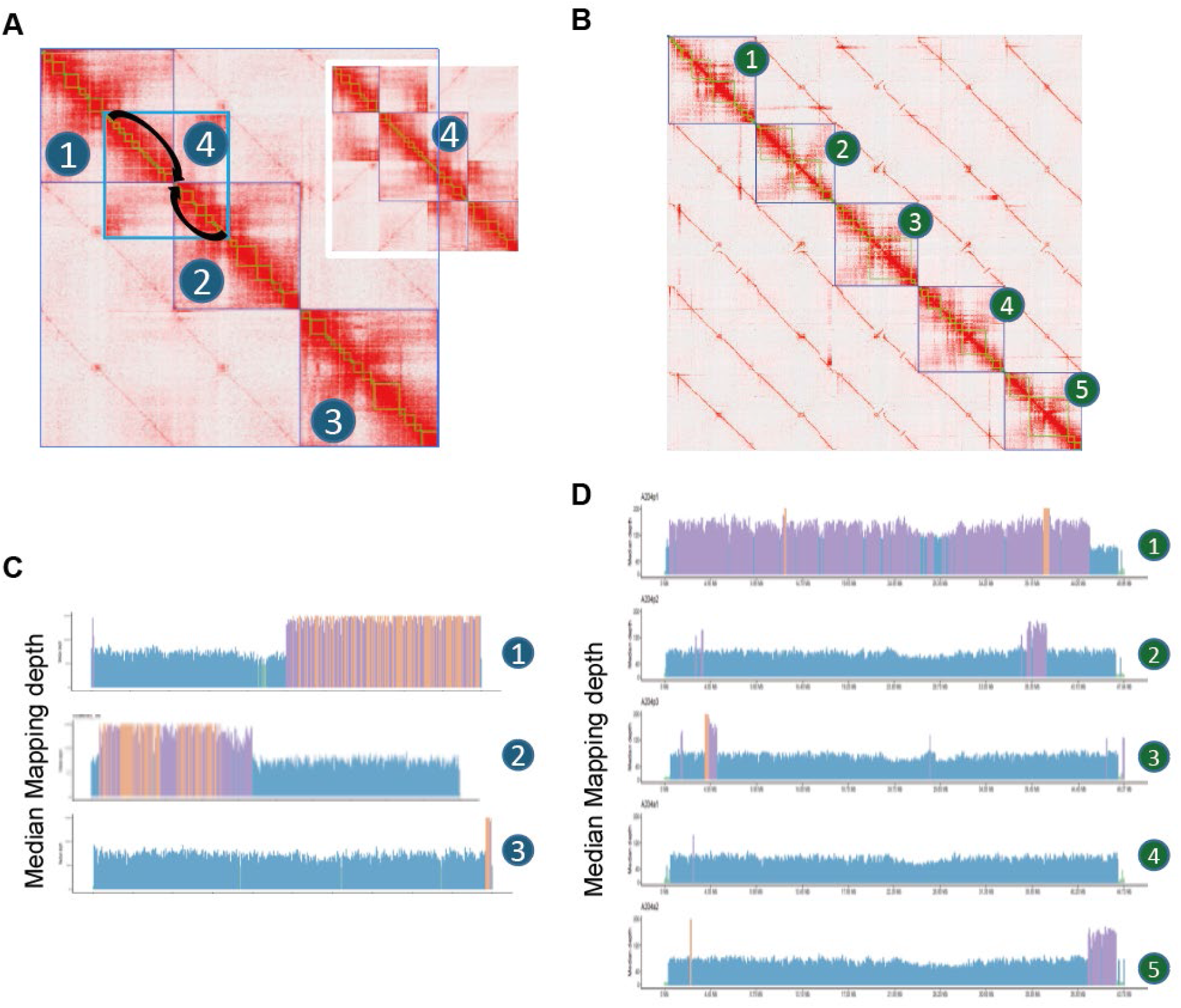
Hi-C interaction and read-depth patterns identify collapsed regions in the *Brachypodium phoenicoides* Bpho6 and *B. boissieri* Bbois3 assemblies. (A, B) Hi-C contact matrices for representative collapsed regions in the Bpho6 (A) and Bbois3 (B) genome assemblies. The blue boxes in panel A indicate genomic regions with abnormal interaction patterns consistent with collapse. (C, D) The Illumina short-read mapping depth across the corresponding chromosome (marked with the same number) in Bpho6 (C) and Bbois3 (D), summarized as the median depth in non-overlapping 100-kb windows. Windows are color-coded by their depth class relative to the bimodal depth distributions in Fig. S2: blue marks windows whose depths fall within the first peak (non-collapsed regions; e.g., 37-109 for Bpho6), whereas purple and yellow mark windows whose depths fall within the second peak (collapsed regions; e.g., 110-200 for Bpho6). Yellow indicates windows whose median depth exceeds the plotting maximum and is therefore truncated.

To recover collapsed regions between haplotypes/subgenomes, we designed our CollapsedChrom pipeline (Fig. 3) for detecting and rescuing collapsed segments using the C-Phasing collapse module, guided by our prior knowledge of the expected copy number of each haplotype/subgenome (see Methods). In Bpho6, we rescued a total of 407 collapsed regions, recovering 328,915,430 bp of sequence. In Bbois3, we rescued 112 collapsed regions, with a total of 195,813,346 bp of recovered sequence (Fig. 4). To validate the correct restoration of these regions, we reassigned hifi reads to the collapsed-rescued assemblies and quantified the read depth patterns at the genome level. The depth-frequency distribution showed a single dominant peak, and sliding-window depth-of-read analyses further indicated that previously collapsed regions now exhibit the expected depth (Fig. S3). Notably, the total number of chromosomes in the rescued sets also matched experimental evidence. In addition, we performed whole-genome alignments with the two reference genomes and observed the expected syntenic relationships: Bpho6 to *B. sylvaticum* showed a 4:1 match (subgenomes G and E2), and Bbois3 to *B. retusum* (subgenome A2) showed a 6:1 match (Fig. S4). It should be noted that because the allotetraploid Bpho6 genome consists of two subgenomes with different chromosomal basic number each (G with x=9 and E2 with x=5), and because the E2 x=5 chromosomes probably resulted from four Nested Chromosome Fusions (NCFs) of an intermediate ancestral *Brachypodium* genome with x=9^15,16^, the G x=9 chromosomes of Bpho6 (18 chromosomes) show almost perfect matches in full length with the G x=9 chromosomes of *B. sylvaticum*. In contrast, the remaining E2 x=5 chromosomes of Bpho6 (10 chromosomes) also show high synteny with the G x=9 chromosomes of *B. sylvaticum* but with different rearrangements, corresponding to those four NCFs (Fig. S4). Taken together, these independent lines of evidence support the accuracy and robustness of our collapse detection and rescue strategy and indicate that the final assemblies faithfully capture haplotype/subgenome-specific sequences at the chromosomal level.

**Figure 3.**
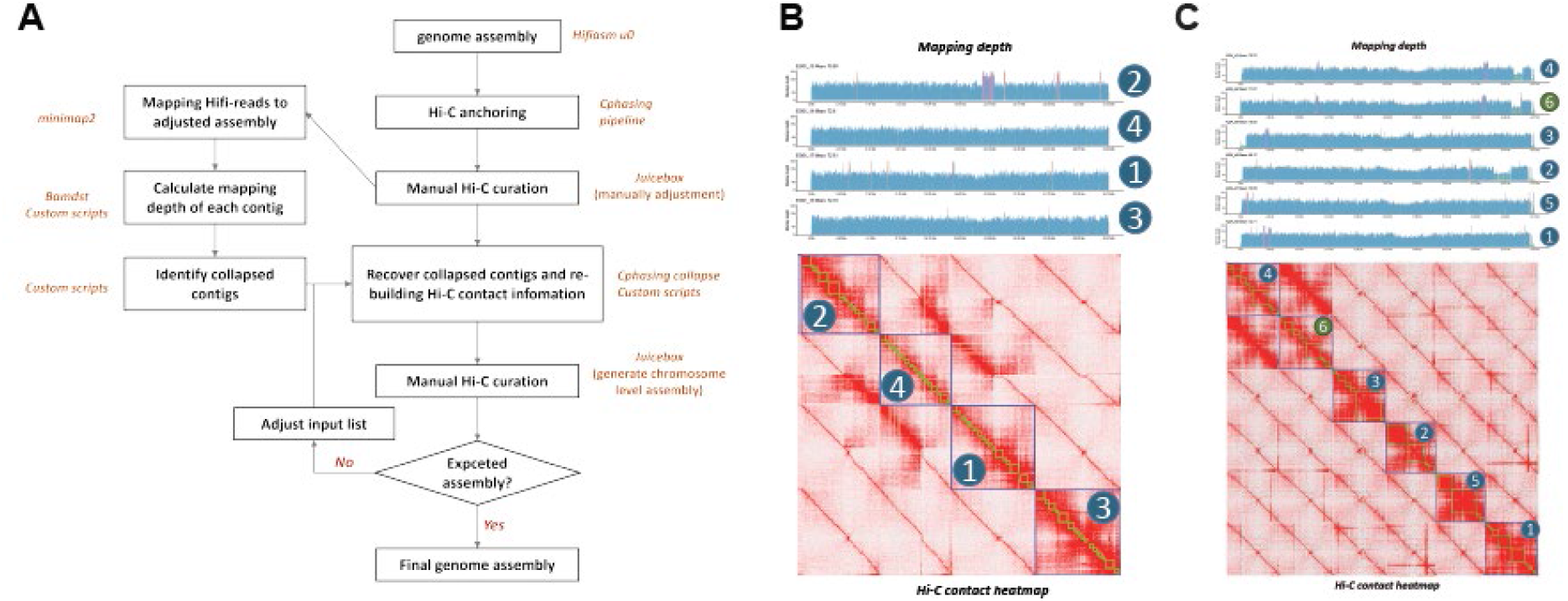
CollapsedChrom pipeline for recovering collapsed assembly segments and representative corrected regions in *Brachypodium phoenicoides* Bpho6 and *B. boissieri* Bbois3. (A) Schematic overview of the collapsed-region recovery pipeline. (B, C) Representative genomic regions after applying the pipeline to Bpho6 (B) and Bbois3 (C), corresponding to the same loci shown in Fig. 2. For each species, the upper tracks show read mapping depth after recovery, and the lower panels show the Hi-C contact heatmaps, illustrating that regions previously exhibiting depth inflation and abnormal interaction patterns are resolved into structures consistent with their expected copy number and local chromatin contact organization.

**Figure 4.**
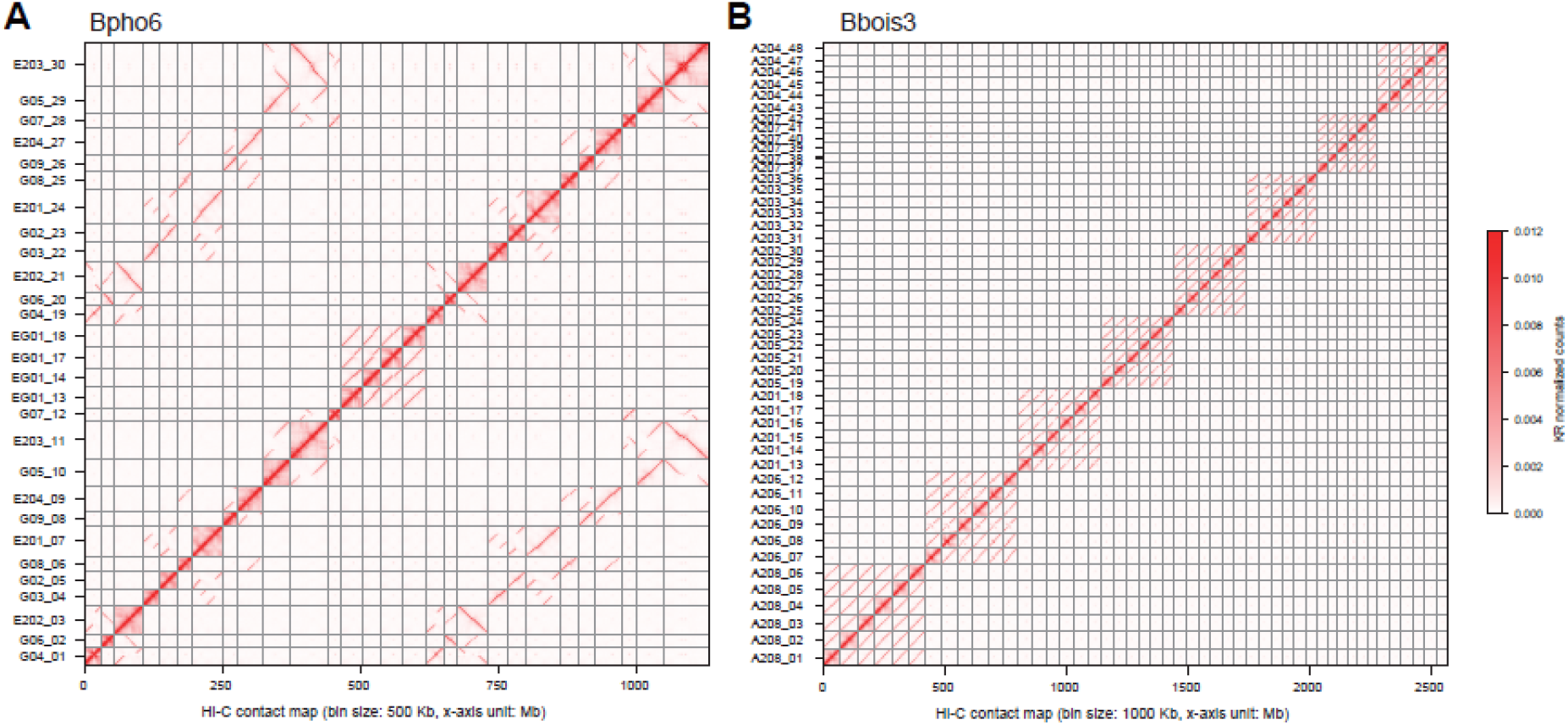
Chromosome-scale Hi-C contact maps of the *Brachypodium phoenicoides* Bpho6 and *B. boissieri* Bbois3 assemblies. Hi-C contact matrix heatmaps for Bpho6 (A) and Bbois3 (B), showing genome-wide chromatin interaction intensity between scaffolds/contigs ordered along the assembled chromosomes (x- and y-axes, Mb). The results highlight well-defined chromosome-scale interaction blocks consistent with the final scaffold structure.

### Chromosome-level genome annotation

Following the targeted recovery of collapsed regions and rigorous manual curation, we generated final chromosome-level assemblies for *B. phoenicoides* (Bpho6) and *B. boissieri* (Bbois3) with total lengths of 1.14 Gb and 2.54 Gb, respectively. Assembly quality was thoroughly validated using multiple independent metrics. K-mer analysis using Merqury yielded QV and k-mer completeness scores of 55.75/98.23% for Bpho6 and 51.12/97.63% for Bbois3.

Furthermore, the structural integrity of repeat-rich genomic regions was corroborated by Long Terminal Repeat (LTR) Assembly Index (LAI) scores of 17.47 and 15.25 for Bpho6 and Bbois3, respectively, both falling within the range of “gold-standard” assemblies. Annotation of repetitive elements using the EDTA pipeline revealed that transposable elements (TEs) and other repeats constitute 44.86% of the Bpho6 genome and 55.88% of the Bbois3 genome. Using an integrative gene prediction workflow, we identified 120,704 protein-coding genes in Bpho6 and 189,694 in Bbois3. High annotation completeness was corroborated by BUSCO scores of 99.8% and 99.9%, respectively. At the monoploid level, the longest haplome spanned 588.91 Mb with 62,289 genes in Bpho6, and 445.97 Mb with 31,849 genes in Bbois3. By recovering these previously collapsed regions, we have produced highly accurate and complete chromosome-level references. These high-quality genomes provide a foundation for future evolutionary and functional analyses.

## Conclusions

In this study, we addressed the challenge of sequence collapse in polyploid genome assembly and generated high-quality, chromosome-level reference genomes for the allotetraploid *Brachypodium phoenicoides* (Bpho6) and the hexaploid *B. boissieri* (Bbois3). By implementing the novel rescue pipeline CollapsedChrom based on prior karyotypic knowledge and read-depth profiles, we successfully recovered 328.9 Mb and 195.8 Mb of previously collapsed sequences, respectively. This strategy effectively corrected the initial copy number deficits observed in the primary assemblies and achieved k-mer completeness greater than 97% for both species. Our final assemblies meet gold-standard LAI criteria and provide insight into the complex genomic architecture in these polyploids. These results provide a robust genomic foundation for future research into the evolutionary dynamics and functional biology of the genus *Brachypodium*.

## Methods

### Plant material, library construction and genome sequencing

Seeds of *Brachypodium phoenicoides* (Bpho6) and *B. boissieri* (Bbois3) were collected in the field from native Spanish localities (Bpho6: Spain: Huesca: Panzano, Sierra Guara, 42º12’42.30’’N 0º11’16.62’’W, 625m, marl loams, *Quercus subpyrenaica* opens, 11/10/2016, Leg. P. Catalán; Bbois3: Spain: Granada: Sierra de Huetor, Puerto de La Mora, 30SED50 X461676 Y4123355, 05/08/2011, Leg. E. Salmerón). Seeds from each individual were kept separately, labeled, germinated, and grown in a controlled growth chamber under 14 h light / 10 h dark cycles at 23 ± 2 °C, using Pindstrup Substrate (Pindstrup Mosebrug, Pindstrup, Denmark; www.pindstrup.dk). Fresh leaf tissue was collected from a single plant per accession for genome and transcriptome sequencing, including PacBio HiFi, Illumina short reads, Hi-C, and RNA-seq. For PacBio HiFi sequencing, SMRTbell libraries were prepared following the manufacturer’s protocol (Pacific Biosciences, CA, USA) with a 15 kb size-selection strategy. Sequencing was performed on a PacBio Revio platform. Illumina paired-end libraries were constructed from the same plant and sequenced on an Illumina NovaSeq 6000 platform. For Hi-C library preparation, fresh leaf tissue was cross-linked with 1% formaldehyde, and the chromatin was digested with the restriction enzyme DpnII. Proximity-ligated DNA fragments were processed following standard Hi-C library preparation procedures and sequenced on an Illumina NovaSeq 6000 platform. Total RNA was extracted from fresh leaves using the RNeasy Plant RNA Kit (Qiagen), and RNA-seq libraries were prepared using standard protocols and sequenced in paired-end mode on an Illumina NovaSeq 6000 platform. Raw Hi-C reads and short Illumina reads were first processed with fastp^31^ v2.0 using the parameters -q 20 -5 -3 -w 10 to trim and filter low-quality reads.

### Genome survey

K-mer depth distributions were computed from the HiFi reads using FastK (https://github.com/thegenemyers/FASTK) with k = 31. The resulting k-mer spectra were modeled with GenomeScope^29^ v2.0 to infer key genomic properties, such as haploid genome size and heterozygosity. Finally, we inferred genome ploidy using Smudgeplot^29^ v0.5.0, which leverages FastK-derived k-mer databases and heterozygous k-mer pair coverage patterns (“het-mers”) to infer ploidy from the sequencing data.

### Genome assembly and quality estimation

We performed Hi-C integrated genome assembly using hifiasm^30^ v0.25.0-r726 (hifiasm improved assembly quality since v0.24). Aside from specifying -u 0 and -t 60, all other parameters were left at their default values. Inspection of the hifiasm log files indicated homozygous read coverage values of 149 for Bpho6 and 156 for Bbois3, consistent with the k-mer–based genome survey. We also evaluated alternative assemblers, including HiCanu^32^ and Verkko2^33^, but none produced assemblies that outperformed our hifiasm results. Furthermore, we tested multiple combinations of hifiasm parameters, focusing on -D, -r, -a, - s, and -l; however, these adjustments yielded no improvements or only marginal ones.

C-Phasing^23^ v0.2.623 was used to align filtered Hi-C reads with their corresponding assembled contigs and perform chromosomal scaffolding, which clusters, orders, and orients the contigs using Hi-C interaction information Subsequently, the Hi-C contact matrices were visualized and manually processed in Juicebox v1.11.08 to correct residual disassemblies and refine the scaffold structure.

Assembly quality was evaluated using Merqury^34^ v1.3 based on k-mer derived metrics. Additionally, filtered Illumina short reads from Bpho6 and Bbois3 were mapped to their corresponding assemblies using BWA-MEM2^35^ v2.2.1, and the resulting alignments were processed into sorted BAM files. SAMtools^36^ v1.21 and Bamdst (https://github.com/shiquan/bamdst) were used to summarize mapping rate, coverage, and read-depth distributions for each assembly. The LTR Assembly Index (LAI)^37^ values were computed with LTR_retriever^38^ v2.9.0 using default parameters.

### Recovery of collapsed regions

To improve accuracy at the chromosomal scale, we implemented a collapsed-region recovery pipeline CollapsedChrom (https://github.com/Bioflora/Collapsed-chromosomal-assemblies-pipeline) that integrates mapping-based detection with Hi-C-guided iterative scaffolding using C-Phasing^23^ v0.2.623. First, we mapped sequencing reads to the initial assembly unitigs using minimap2^39^ v2.21 with parameters --secondary=no -x map-hifi -a. Next, we calculated window-based sequencing depth using bamdst. Unitigs larger than 20 kb, for which >30% of the total length showed high depth relative to the assembly baseline, were considered candidates for collapsed segments.

Candidate collapsed unitigs were then processed using custom scripts (available at https://github.com/Bioflora/Collapsed-chromosomal-assemblies-pipeline) to duplicate underrepresented segments and thus reconstruct the missing homologous copies. In parallel, we updated the original Hi-C matrix by adding interaction information for the newly generated copies, so that downstream scaffolding could fully exploit Hi-C linkage patterns. All duplicated fragments were consistently renamed and tracked to preserve provenance and to facilitate anchoring into the correct homologous groups during subsequent Hi-C– based clustering.

After recovering the collapsed region, we applied C-Phasing’s collapse and scaffolding modules to reconstruct the Hi-C contact matrices and generate chromosome-scale scaffolds. The resulting contact matrices were inspected and manually processed in Juicebox to correct residual misjoins and refine scaffold structure. We then remapped sequencing reads to the recovered assembly and re-evaluated mapping depth. We also performed a whole-genome alignment between the recovered assembly and the genome of its closest relative to further validate the assembly. If residual collapse was still detected (for example, a region expected to be present in four copies but assembled as one), we repeated the same duplication procedure and performed another round of read mapping and depth inspection. This iteration continued until all recoverable collapsed regions were resolved.

### Genome annotation

EDTA^40^ v2.2.2 was used to identify and annotate repetitive elements in the Bpho6 and Bbois3 genomes. We predicted gene models using three complementary strategies: ab initio, homology-based, and transcriptome-based annotation. For ab initio prediction, we employed ANNEVO v2.2 (https://github.com/xjtu-omics/ANNEVO), Helixer^41^ v0.3.6, and Eviann^42^ v2.0.4. For homology-based prediction, GeMoMa^43^ v1.9 was run using protein sequences from *Brachypodium distachyon* (Bd21, https://phytozome-next.jgi.doe.gov/info/Bdistachyon_v3_1), *B. stacei* (Bsta ECI^44^ and Bsta ABR114, https://phytozome-next.jgi.doe.gov/info/Bstacei_v1_1), *B. hybridum* (Bhyb ECI^13,44^, Bhyb26^45^, and Bhyb ABR113, https://phytozome-next.jgi.doe.gov/info/Bhybridum_v1_1), *B. sylvaticum*^14^, and B. *mexicanum* (https://phytozome-next.jgi.doe.gov/info/Bmexicanum_v2_1), *B. arbuscula* (https://phytozome-next.jgi.doe.gov/info/BarbusculaBARB1_v3_1). For transcriptome-based prediction, RNA-seq reads were aligned to the genome with HISAT2^46^ v2.2.1, and coding sequences were inferred from transcript assemblies using StringTie^47^ v3.0 and TransDecoder v5.7.1 (https://github.com/TransDecoder/TransDecoder). Finally, gene models from all approaches were integrated into a non-redundant consensus set using EvidenceModeler^48^ (EVM) v1.1.1, and annotation completeness was assessed with BUSCO^49^ v5.8.3.

## Data availability

The custom computational pipeline developed for the detection and rescue of chimeric and collapsed genomic regions in this study is freely available on GitHub at https://github.com/Bioflora/Collapsed-chromosomal-assemblies-pipeline. The raw sequencing data generated and analyzed during the current study are available from the corresponding author upon reasonable request and subject to appropriate permissions.

## Acknowledgements

We thank researchers of the Bioflora group at High Polytechnic School of Huesca - University of Zaragoza (EPSHU-Unizar) for the propagation of the Bpho6 and Bbois3 seeds, researchers at the Joint Genome Institute (JGI) for other *Brachypodium* reference genomes deposited in Phytozome, and we thank them and Drs. Joanna Lusinska and Robert Hasterok from the University of Silesia in Katowice for the fruitful discussions on the karyotypes, genomic composition, and evolution of the studied *Brachypodium* polyploids, and the technicians and students from Lanzhou University for their assistance with DNA and RNA isolation. Genomic sequencing procedures were performed at Xi’An Haorui Gene Technologies Ltd. (China). Genomic assembly and annotation analyses were carried out at the High Polytechnic School of Huesca, University of Zaragoza (Spain).

## Contributions

WM assembled and annotated the *Brachypodium phoenicoides* Bpho6 and *B. boissieri* Bbois3 reference genomes, developed the CollapsedChrom pipeline, and drafted the preliminary manuscript. JL and PC supervised the genomic analyses of Bpho6 and Bbois3 and wrote the manuscript. The authors declare no conflict of interest.

---

**Figure S1.**
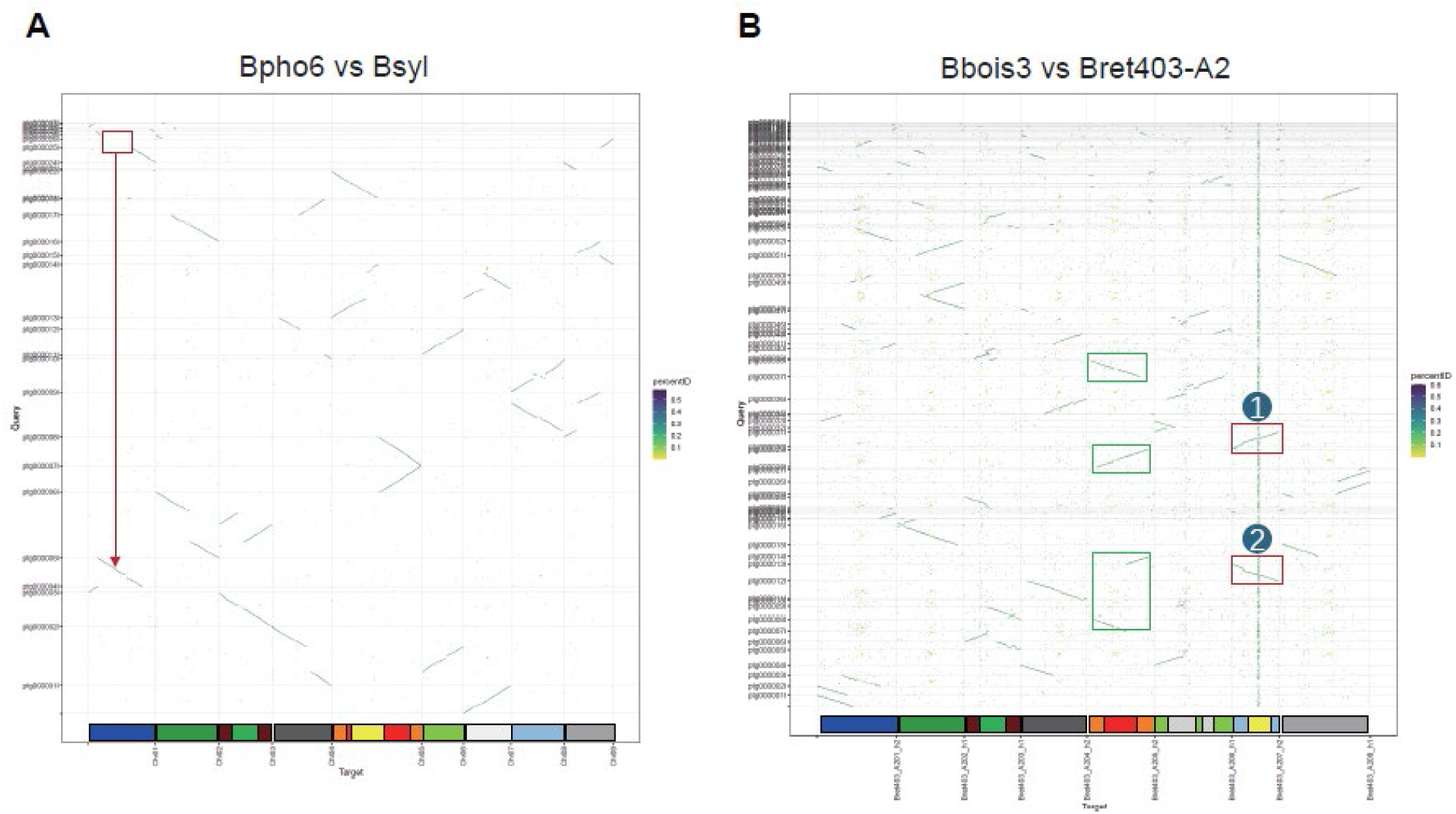
Whole-genome synteny dot plots reveal missing homologous segments in the initial primary assemblies of *Brachypodium phoenicoides* Bpho6 and *B. boissieri* Bbois3. (A) Genome-wide alignment dot plot between the initial primary assembly of Bpho6 and the closely related reference genome B. sylvaticum (Bsyl-Ain1). (B) Genome-wide alignment dot plot between the initial primary assembly of Bbois3 and the closely related reference genome Bret403-A2. Points represent homologous alignments (colored by sequence identity). Red boxes highlight regions that are present in the reference but absent from the corresponding initial primary assembly, indicating missing/collapsed segments, whereas green boxes mark regions with well-aligned synteny. Karyotype barplots shown below each panel follow Sancho et al. (2022).

**Figure S2.**
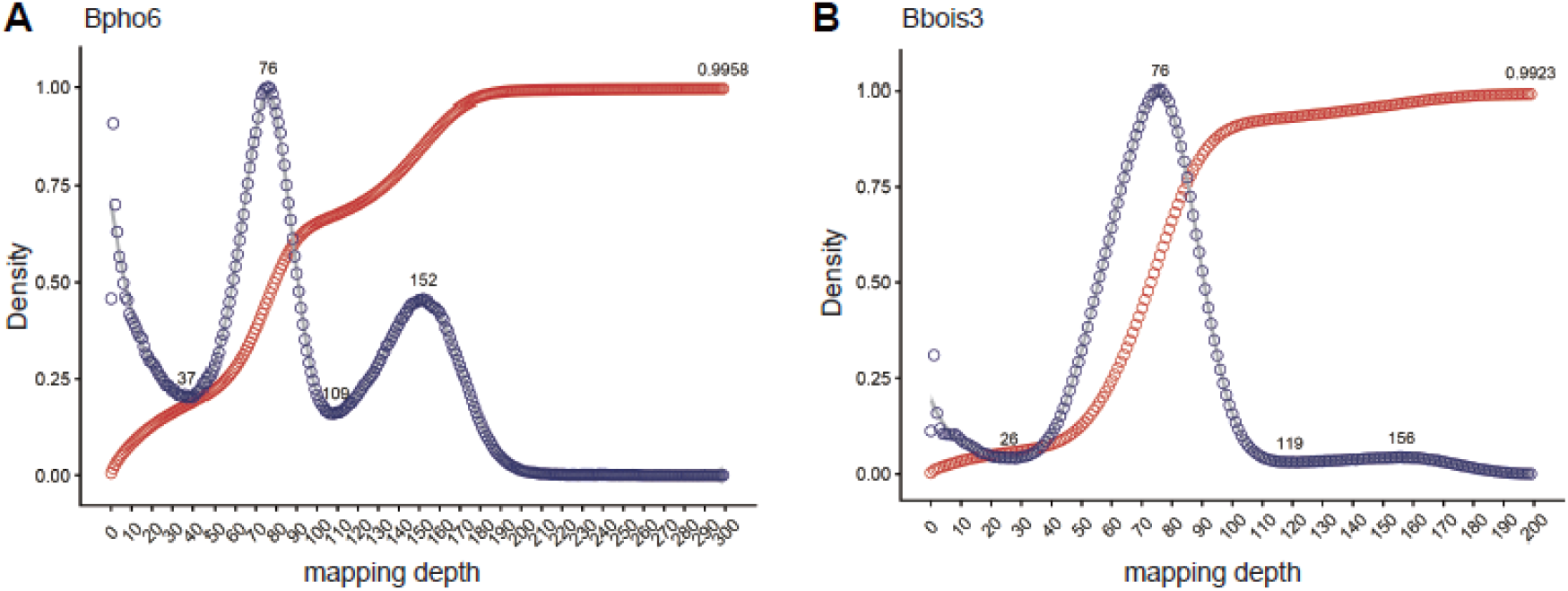
HiFi read mapping depth distributions in the pre-recovery assemblies of *Brachypodium phoenicoides* Bpho6 and *B. boissieri* Bbois3. (A, B) Distribution of HiFi read mapping depth (blue) and the corresponding data cumulative proportion curve (red) after mapping HiFi reads to the unitig-based assemblies following C-Phasing scaffolding and manual curation (i.e., before collapsed-region recovery) for Bpho6 (A) and Bbois3 (B).

**Figure S3.**
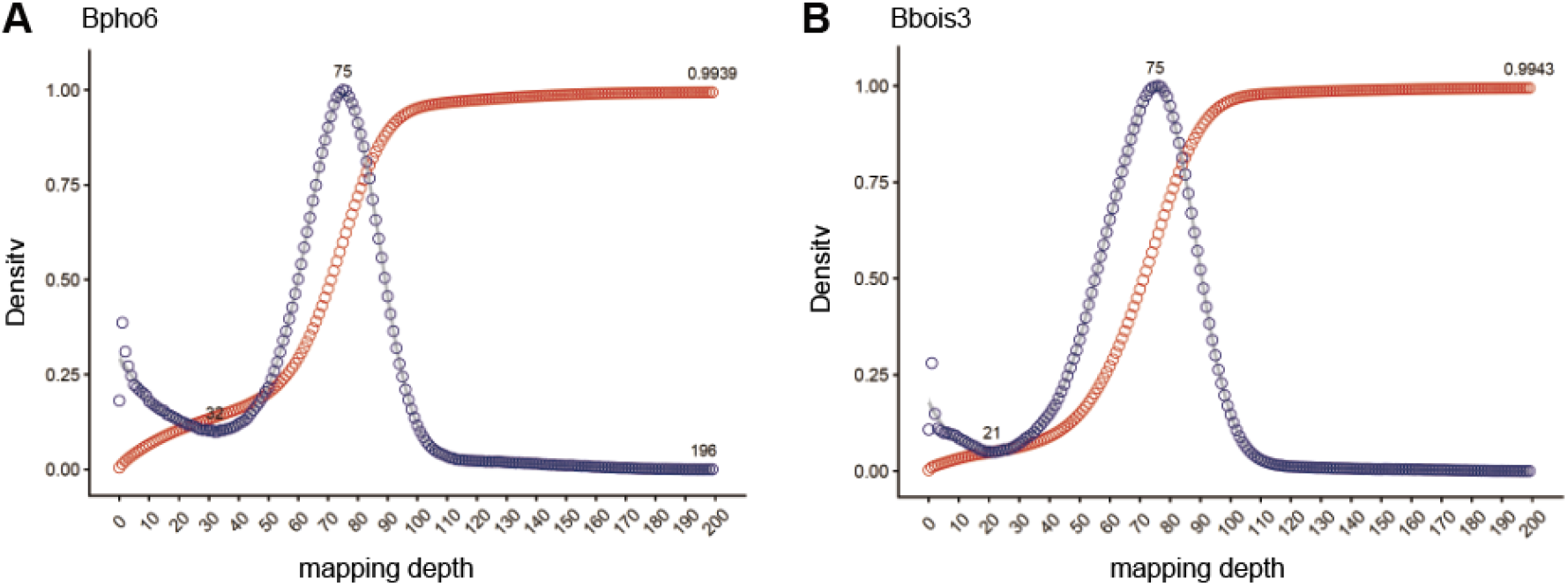
HiFi read mapping depth distributions in the post-recovery assemblies of *Brachypodium phoenicoides* Bpho6 and *B. boissieri* Bbois3. (A, B) Distribution of HiFi read mapping depth (blue) and the corresponding data cumulative proportion curve (red) after mapping HiFi reads to the assemblies following pipeline showed in Fig. 3

**Figure S4.**
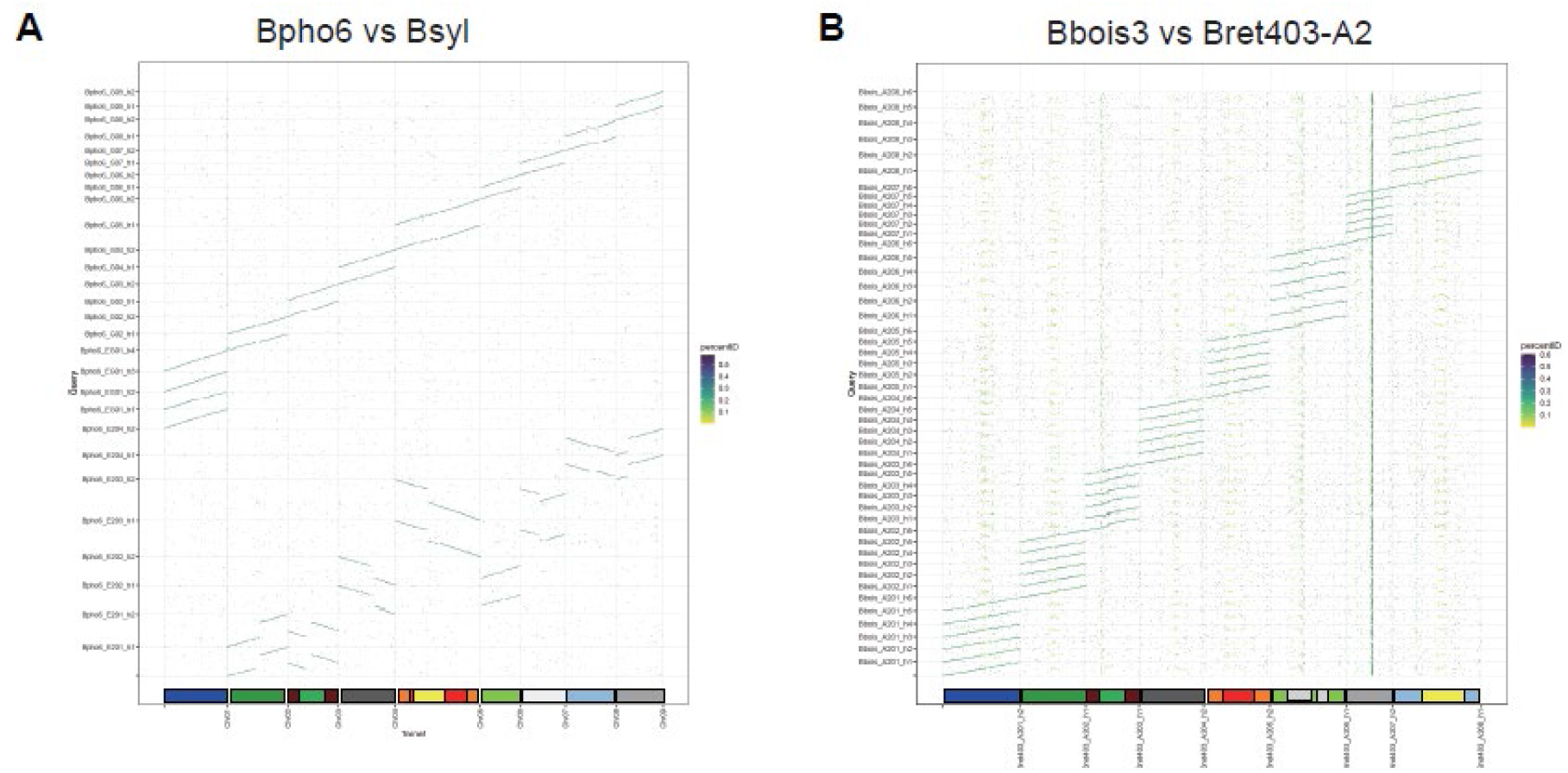
Whole-genome synteny dot plots after collapsed-region recovery highlight restored homologous segments in *Brachypodium phoenicoides* Bpho6 and *B. boissieri* Bbois3. (A) Genome-wide alignment dot plot between the post-pipeline Bpho6 assembly and the closely related *B. sylvaticum* (Bsyl) reference genome. (B) Genome-wide alignment dot plot between the post-pipeline Bbois3 assembly and the closely related Bret403-A2 genome. Dots indicate homologous alignments (colored by sequence identity). Karyotype barplots shown below each panel follow Sancho et al. (2022).

